# Optogenetic activation of visual thalamus generates artificial visual percepts

**DOI:** 10.1101/2022.12.07.519434

**Authors:** Jing Wang, Hamid Azimi, Yilei Zhao, Melanie Kaeser, Pilar Vaca Sánchez, Michael Harvey, Gregor Rainer

**Author notes:** To Whom correspondence should be addressed, Prof. Gregor Rainer, Section of Medicine, University of Fribourg, Chemin du Musee 5, CH-1700 Fribourg, Switzerland.

## Abstract

The lateral geniculate nucleus (LGN), a retinotopic relay center where visual inputs from the retina are processed and relayed to the visual cortex, has been proposed as a potential target for artificial vision. At present, it is unknown whether optogenetic LGN stimulation is sufficient to elicit behaviorally relevant percepts and the properties of LGN neural responses relevant for artificial vision have not been thoroughly characterized. Here, we demonstrate that tree shrews pretrained on a visual detection task can detect optogenetic LGN activation using an AAV2-CamKIIα-ChR2 construct and readily generalize from visual to optogenetic detection. We also characterize how amplitude and frequency of optogenetic stimulation affect behavioral performance. Given the importance of temporal stimulation parameters, we also study tree shrew behavioral ability to discriminate among pairs of visual flicker frequencies and demonstrate performance transfer among related discrimination problems as well as limitations. Neural recordings in LGN and primary visual cortex (V1) examining two flicker discriminations reveal that while LGN neurons multiplex information about temporal stimulation discriminations, V1 neurons explicitly encode information about one or the other discriminations but not both. Our findings confirm the LGN as a potential target for visual prosthetics and introduce a distinction in stimulus encoding between thalamic and cortical representations that could be of fundamental importance in artificial vision.

## Introduction

Vision loss is widespread and rapidly progressing worldwide due to an aging population, with over 45 million people suffering from blindness in 2020 and over 55 million cases forecast for 2050 ^1^. For many eye diseases, including glaucoma, macular degeneration and retinitis pigmentosa there are no available treatments, and they progressively lead to complete loss of vision ^2^. It has been shown that prosthetic electrical stimulation in visual structures can produce visual percepts, in the form of phosphenes, and at least partially restore visual function ^3^. To date, the most advanced of these approaches are retinal implants, with sub- and epiretinal approaches that have both been extensively tested in human patients. Despite considerable progress, the percepts generated by these prostheses remain far from natural vision, and many patients stop using their implants and often report limited improvements in quality of life ^4^. It is interesting to note that phosphenes tend to vary substantially in terms of their appearance between patients, but they do not change much with time ^5^. This suggests that differences in percept may result not from neural plasticity but rather from variations in device implantation such that slightly different parts of the retinal circuitry are activated by the implant in each individual. Indeed, the retina is a highly complex computational engine containing dozens of distinct, specialized cell types ^6^, and electrical stimulation is not cell-type specific and can trigger activation in any neurons within a defined volume around the electrode tip. Another pertinent limitation relates to the size of the retinal implants, which can cover only up to about 20 degrees of visual angle such that patients must make head movements to repeatedly scan the environment during locomotion to detect obstacles, walls or doors. The interpretation of phosphenes generated by retinal prosthetic devices generally does not becomes effortless or intuitive but continues to require considerable cognitive resources and effort. Nevertheless, technical advances under development ^7^ may certainly enhance the usefulness of retinal implants in the future.

The primary visual cortex (V1) has also been investigated as a target for electrical visual prosthetics, due to attractive features such as its large surface area, could enable highly precise representations of the visual environment using suitable stimulation arrays. Early work in this area has demonstrated proof-of principle ^8,9^ that electrical activation of V1 can elicit phosphene percepts, and there has been resurgent interest in cortical visual prostheses in recent years ^10,11^. For example, it was recently shown that monkeys could detect, and make saccades to, V1 stimulation at particular retinotopic sites, as well as interpret multi-electrode stimulation in the context of a learned visuomotor association task ^12^. This work relied on the implantation of over 1000 penetrating electrodes in multiple arrays in visual cortex, highlighting the technical challenge of broad coverage for optimizing visual prosthetic stimulation. In another recent study, blind human subjects readily identified sequential electrical activation on a rectangular grid of flexible electrodes spaced 1cm apart that represented letter symbols, whereas simultaneous electrical activation was much harder to comprehend ^13^. This latter finding suggests more generally that sequential patterned activation across multiple sites could be highly useful in producing more coherent percepts. Numerous issues remain however in relation to cortical electrical visual prostheses, including problematic surgical access to the large part of V1 that is buried in the calcarine sulcus particularly for penetrating electrodes, potential damage caused by the electrodes during implantation, long term stability of the electrode-neural interface, and risk of triggering seizures due to the highly recurrently connected nature of the cortex. An advantage of V1 prosthetics is that they are more general and can be applied also in eye disorders where the retinal ganglion cells are damaged, such as glaucoma. This same advantage would also apply to the lateral geniculate nucleus (LGN) of the thalamus, where the ganglion cell axons synapse onto the thalamocortical neurons whose axons deliver visual information to cortex ^14,15^. Indeed, electrical activation of LGN has been shown to activate V1 neurons ^16^, and LGN activation can be detected by monkeys, who are able to saccade reliably to the corresponding visual field location ^17^. In terms of prosthetics, the LGN has another advantage in that it is a six layered retinotopically organized nucleus where information pathways are segregated and could therefore be specifically targeted ^18,19^. However, the LGN is a deep brain structure far from the cortical surface and has to date not been much studied in terms of prosthetic applications.

Gene therapy and optogenetics have also brought novel approaches for combating eye diseases. For example, the light sensitive channel-rhodopsin protein was expressed in retinal ganglion cells of a patient suffering from photoreceptor loss due to retinitis pigmentosa, leading to recovery of some visual function using suitable illumination via goggles ^20^. Optogenetics also opens another avenue for vision restoration, combining viral delivery to activate elements of the visual pathway with patterned laser or LED light stimulation to produce visual percepts ^21^. The cell-type specificity of optogenetics is likely to confer advantages compared to electrical stimulation, as only cell-types of interest can be activated. Both cortex and LGN for example contain numerous inhibitory interneurons, whose activation may hamper the efficacy of prosthetic stimulation.

In the present study, we focus on the LGN as a potential target of optogenetic restoration of vision using optogenetic activation of thalamocortical projecting neurons. We employ ChR2 coupled to the CamKIIα promotor, as it is known to induce activation as well as plasticity in excitatory neurons ^22,23^. We characterize opsin expression in LGN and V1 and assess the impact of opsin activation on both LGN circuit activity, and behavioral detection task performance linked to optogenetic activation. We use tree shrews as an animal model in this study, as they are a diurnal mammalian species closely related to primates and humans that have multiple advantages for the translational study of visual prosthetics. For example, tree shrew V1 has a large surface area of 4000 mm^2^ and exhibits orientation columns and shares many other aspects of functional organization with primates and humans ^25–28^. Their retina is composed largely of cones ^29^, which is useful for prosthetic work as humans tend to illuminate their environments; they are a diurnal mammal that is highly reliant on vision ^30^ and can readily be trained on cognitive visually based behavioral tasks ^31,32^. In this study, we also assess behavioral and neuronal processing of temporally modulated visual stimuli, and compare it to optogenetically triggered activation, with the goal to better understand similarities and differences of LGN and V1 activations to natural vision and artificial optogenetic activation.

## Results

We injected adeno-associated (AAV2) virus that contained a construct for the light-sensitive ion channel ChR2 and the enhanced red fluorescent protein mCherry under the control of the CamKIIα promotor into the tree shrew LGN. In post-mortem analyses, we performed immunohistochemistry on the LGN sections for parvalbumin (PV), which revealed the characteristic laminar organization of the LGN (Fig.1A). We observed somewhat less prominent expression in LGN layer 3, which is consistent with a previous report documenting paler parvalbumin staining in layers 3 and 6 ^33,34^. We found robust expression of CamKIIα-ChR2 within LGN layers in the vicinity of the injection site including on cell bodies, targeting inner LGN layers 1,2 and 3 (Fig.1A). As the pattern of PV expression in particular cell types varies between species ^35^, we examined co-expression of PV with CamKIIα-ChR2 for our dataset. We found that 29% of mCherry labelled CamKIIα neurons also expressed PV (Fig.1B), suggesting that the CamKIIα promotor targeted both PV and non-PV expressing excitatory, putative thalamocortical relay cells ^36^. The clear labelling of thalamocortical axons observed in layers 3 and 4 of V1 confirms this (Fig.1C). Both cortical recipient layers contain axons expressing CamKIIα-ChR2, which highlights that neurons in both LGN layers groups 3/6 as well as 1/2/4/5 robustly express the light-sensitive ion channel, as these project respectively to V1 layers 3 and 4. Note that in macaques, CamKIIα is expressed mostly in intralaminar LGN layer, konio-type relay neurons ^37,38^. For functional validation, we performed terminal experiments under isoflurane anesthesia using optrodes that allowed simultaneous recording of neural activity and activation of ChR2 using an optic fiber within the LGN. Results for an example recording site are shown in Fig.2A, illustrating entrainment of LGN spiking activity to light transients at several frequencies of stimulation. Frequency analysis of the evoked spike trains reveals clear peaks at the stimulation frequency, as well as harmonics. Note that at the 80Hz stimulation frequency, the neural response adapts rapidly, and the neuron tends to respond only to the onset of stimulation; a finding that is expected due to ChR2 channel dynamics but indeed also resembles the activation profile to high frequency visual flicker ^27^. The LGN can be activated in tonic or burst mode which favor either stimulus fidelity or detectability respectively, ^39^. Action potentials in both cases, however, convey stimulus-specific information to the cortex ^40,41^. We therefore examined to what degree the optogenetic activation triggered bursts in our recordings. For the example neuron (Fig.2A), bursts were rarely triggered and burst percentage of total spikes amounted to 0.7±0.3%, with other recorded neurons also in this range (n=7, range: 0.03% to 3.0%). Optogenetic LGN activation thus elicited mostly tonic spikes, consistent with reports of a moderate propensity for bursting in tree shrew LGN ^42^. Tree shrew LGN layers are specialized for ON- and OFF-type visual inputs ^18^, responding to onset of bright and dark targets with respect to background illumination respectively, as is also the case for other species such as the macaque ^43^. Local recordings of LGN neural activity at the site of light stimulation can thus be used to functionally identify the stimulated layer, as illustrated for 1Hz and 5Hz full-field bright visual stimulation (Fig.2B). Together, these results suggest robust expression of ChR2 in thalamocortical relay cells across tree shrew LGN layers under the CamKIIα promotor.

**Figure 1.**
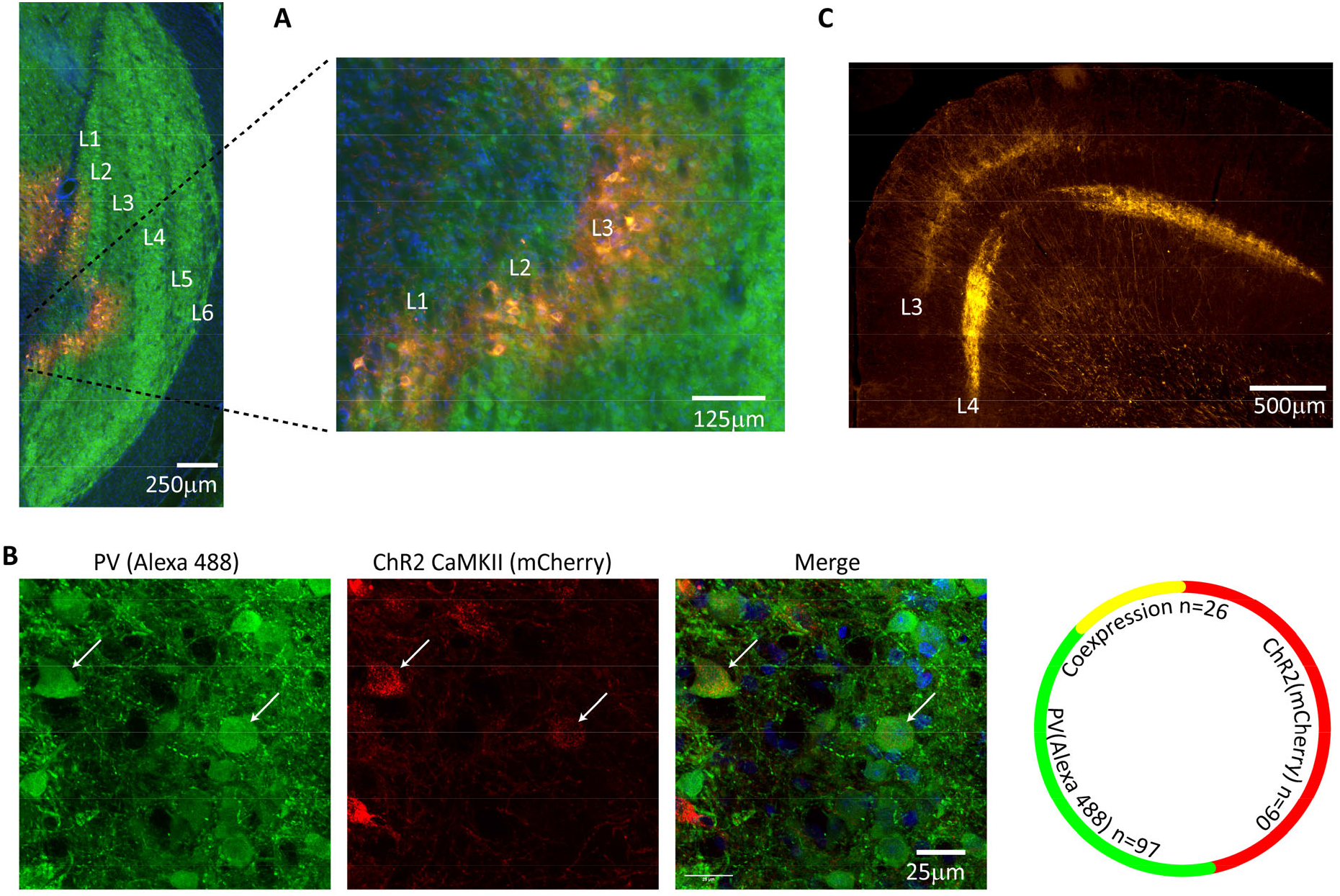
Validation of CaMKIIChR2a expression in Tree Shrew LGN. (A) Fluorescence image of Tree Shrew LGN immunostained for Parvalbumin, green, revealing the LGN layers, and showing viral expression (mCherry, red) in LGN layers 1-3. (B) At left are confocal images showing the Co-localization of ChR2-mCherry with PV-Alexa 488. At right is the quantification. Note that of the 90 mCherry cells counted, 26, or 29%, also expressed PV. (C) Axonal projection patterns in VI from viral transfected cells in the LGN. Note prominent projections to both granular and superficial layers.

**Figure 2.**
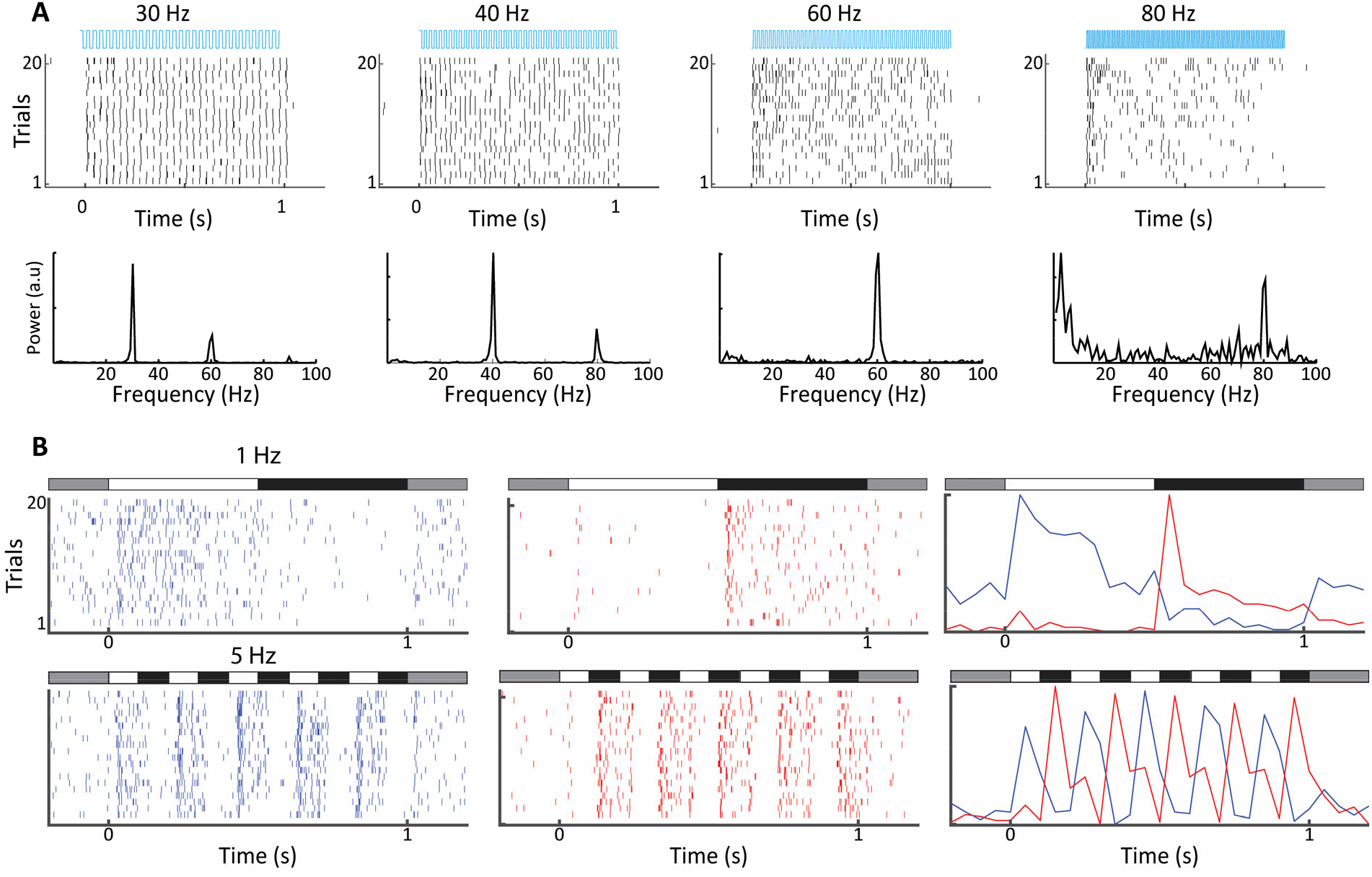
Electrophysiological validation of functional ChR2 expression in LGN CaMKIla neurons. (A) Raster plots, top, show 20 trials of responses of a single LGN neuron to different frequencies of 473 nm blue laser activation. Each vertical bar represents a single spike. FFTs of the same spike trains are shown at the bottom. For this cell phase locking disappeared at the highest frequency, 80 Hz. (B) Raster plots, left and center, and PSTHs, right, reveal both on and off type responses at sites nearby to laser stimulation for both 1Hz top and 5 Hz bottom, stimulation frequencies. Bars at the top of each plot indicate the contrast condition, with grey being the background illumination.

To examine whether animals could detect LGN optogenetic activation, we first trained them on a visual detection task, where they had to enter a nose poke on a transparent horizontal platform at the center of a 70cm diameter opaque sphere and respond to the onset of a visual stimulus by withdrawing from the nose poke (see methods, Fig.3A). Animals were thus in a defined position before and during visual target presentation, allowing us to assess their visual capacities in different parts of the visual field without the requirement of head fixation. We used a 2° phosphene-like stimulus composed of moving white dots to facilitate the subsequent transition to the optogenetic stimulation detection task. Although artificial percepts can take on a variety of forms, they correspond better to moving clouds of dots rather than oriented bars or grating stimuli. Behavioral data for an example tree shrew are shown in Fig.3B for a daily session consisting of 200 trials. It is evident that the animals’ response times are narrow distributed and largely follow the target onset time marked by the thick oblique line with a median response time of 290ms and an overall correct performance of 76%. Our task did not contain any signal absent trials to avoid compromising animals’ motivation for task performance, but we nevertheless used signal detection theory to estimate sensitivity and bias (see methods). Briefly, trials where animals responded within 500ms of target onset were designated as “hits”, whereas responses made during an identical time window where no targets occurred were assigned as “false alarms”, with corresponding “miss” and “correct rejection” assignments. The example dataset in Fig.3B yields a d’ sensitivity value of 2.6. The hit, miss and d’ values during the learning phase for this animal are shown in Fig.3C, illustrating an increasing hit rates as well as decreasing false alarm rates during the seven-day learning period. Group data for the five tree shrews participating in this experiment is shown in Fig.3D, illustrating that all animals showed improved d’ sensitivity over the course of learning, reaching d’ values between 1 and 2.5 at the end of training. Note that the same color is maintained across figures for each animal, allowing comparison of performance of each individual in all the tasks where it participated. The bias for these same behavioral sessions is reveals considerable individual variation as well as a trend for reduced bias with training. Positive and negative bias values signal preponderance of “miss” and “false alarm” type errors respectively. Interestingly, animals with similar and high d’ values of around 2.0, i.e. olive green and bright green colored traces, exhibited opposite bias values suggesting individual differences in strategy and decision threshold. Once animals had acquired the detection task when visual targets were presented at visual field center, we proceeded to study how detection performance generalized to target presentations at other visual field locations. As tree shrews have a wide visual field, we investigated eccentricities of up to 150° in 4 tree shrews, studying a single eccentricity in a single behavioral session. All animals readily generalized detection performance up to 120° (Fig.3E-F), with performance declining at 150°, the highest eccentricity tested, in all except one animal (olive green trace), which detected these peripheral phosphenes without apparent problems. Note that slight differences in head position, estimated at ±15°, cannot be ruled out in the nose poke task, so eccentricity values are not exact as they would be with head fixation. Bias tended to increase with eccentricity (Fig.3G), as animals increasingly missed targets. Taken together, behavioral data on the visual detection task indicates that tree shrews can acquire this task and generalize across the visual field in a freely moving unrestrained behavioral setting, achieving d’ values of up to 2.5 with several weeks of training.

**Figure 3.**
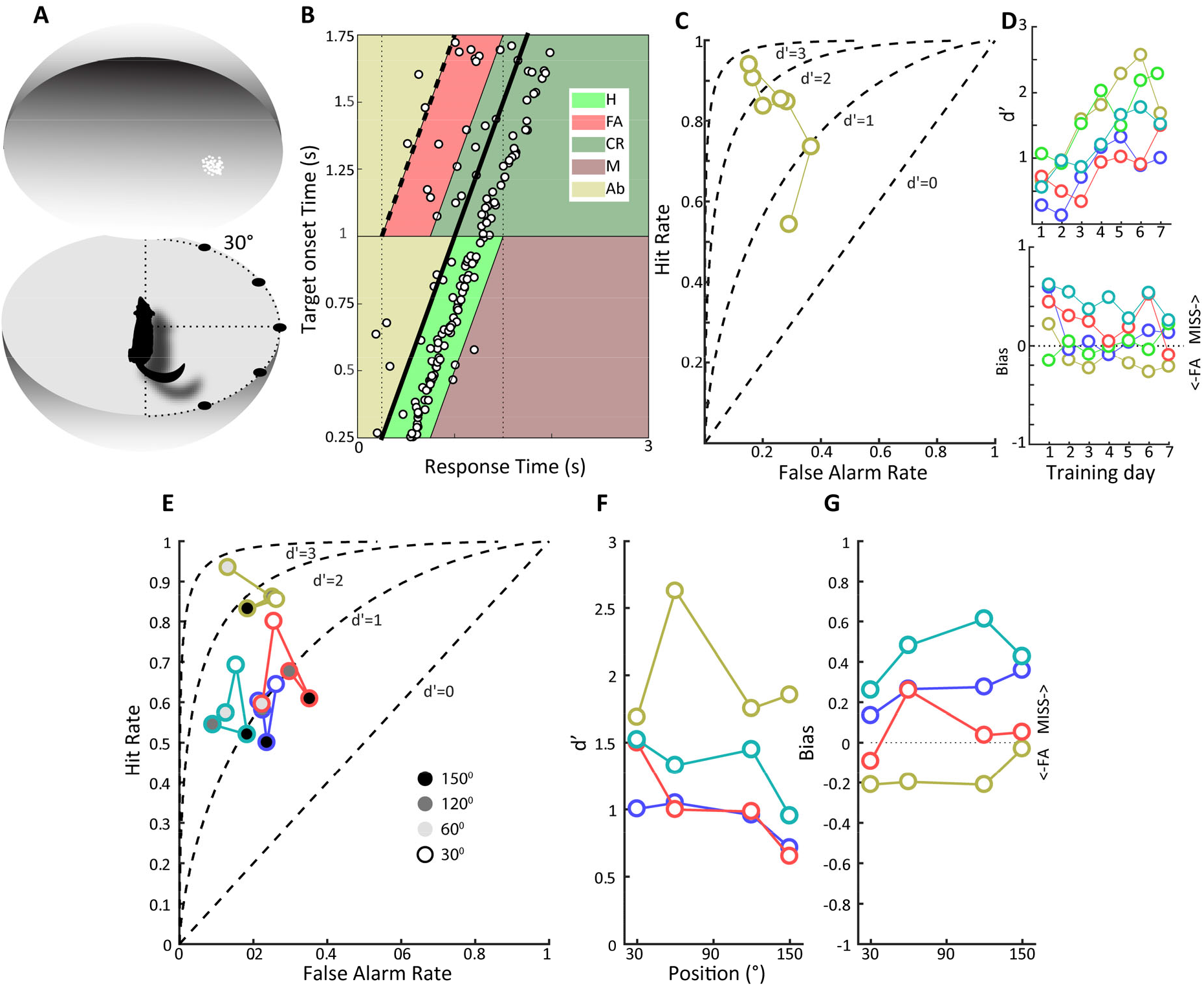
Detection of a visual phosphene. (A) Illustration of the experimental set up. Animals were trained to poke their nose into a response port at the center of a sphere and remain there until the appearance of a 2° patch of moving white dots appeared at the in the field of view. (B) Plotted is the behavior of an animal that has acquired the task when phosphene stimuli were presented at the center of the field of view. Target onset is indicated by the thick, oblique black line. The light green trapezoid depicts the area during which responses were considered hits, and the orange trapezoid depicts the area of false alarms. H, hit, FA, false alarm, CR, correct rejection, M, miss, Ab, abort. (C) False alarm vs hit rates are plotted over training days for the same animal as in B. Note that as learning progresses, the hit rate increases as the false alarm rate declines. (D) At top, d’ values over training days are plotted for all animals, showing increased sensitivity over training days. Different colors denote different animals, and the color scheme is maintained throughout subsequent figures to facilitate comparison of individual animal performance on different tasks. Bottom, bias values for the same animals over training days. Note that bias values below zero indicate more false alarms and those above zero indicate more misses. (E) False alarm vs hit rates are plotted for all animals for phosphene like stimuli presented at different positions relative to the center of the field of view, indicated by grey scale fill. (F) d’ values as a function of stimulus position for the four animals tested. (G) Bias values for the four animals at the different stimulus locations.

We next moved on to study if tree shrews pretrained on visual detection of a phosphene-like stimulus could also detect optogenetic activation in the LGN in the context of the same nose poke task. We tested detection of optogenetic activation using several frequencies (4, 10, 20, 50Hz) and amplitudes (4, 6.5, 10 W) using a single combination of frequency and amplitude in each session, presented in randomized order using a wireless, subdermal radio frequency powered stimulation probe ^44^. An example of behavioral performance during a single session is shown in Fig.4A, corresponding to a behavioral performance of 63% correct trials with a median response time of 260ms and a somewhat broader distribution compared to the visual detection task. Using the same procedure as above, we used signal detection theory to compute sensitivity (d’) and bias for the optogenetic stimulation trials. Results for 3 tree shrews with AAV2 injections and LED implanted in the LGN (olive green, dark blue and bright green traces) and 2 tree shrews injected and implanted in control areas adjacent to the LGN (red and cyan traces) are shown in Fig.4B-C for sensitivity and bias. We note that LGN implanted animals achieved d’ sensitivity values of 1.0 or greater, whereas d’ did not surpass 0.5 in control animals implanted outside the LGN. This indicates that tree shrews can readily detect optogenetic activation of the visual thalamocortical pathway. Importantly, tree shrews immediately generalized from the visual to the optogenetic stimulation, exhibiting d’ values of 0.5, 0.8 and 1.0 respectively on their first session with optogenetic stimulation, suggesting that performance values were not subject to an overall trend but related to specific combinations of amplitude and frequency. Generally, higher amplitudes of stimulation tended to be detected most readily, as would be expected. However, tree shrews differed markedly in terms of which frequency of stimulation was most readily detected, with each of the three animals exhibiting preference for a distinct frequency of stimulation, i.e. 4, 10 and 50Hz for dark blue, bright green and olive green traces. Furthermore, in one animal the preferred frequency differed depending on stimulation amplitude (dark blue trace), with optimum d’ sensitivity at 50Hz for low amplitude and 4Hz for high amplitude, with low sensitivity at the intermediate amplitude. These results highlight a large contribution of individual strategy towards interpreting the optogenetic stimulation, as reported also for humans ^5^.

**Figure 4.**
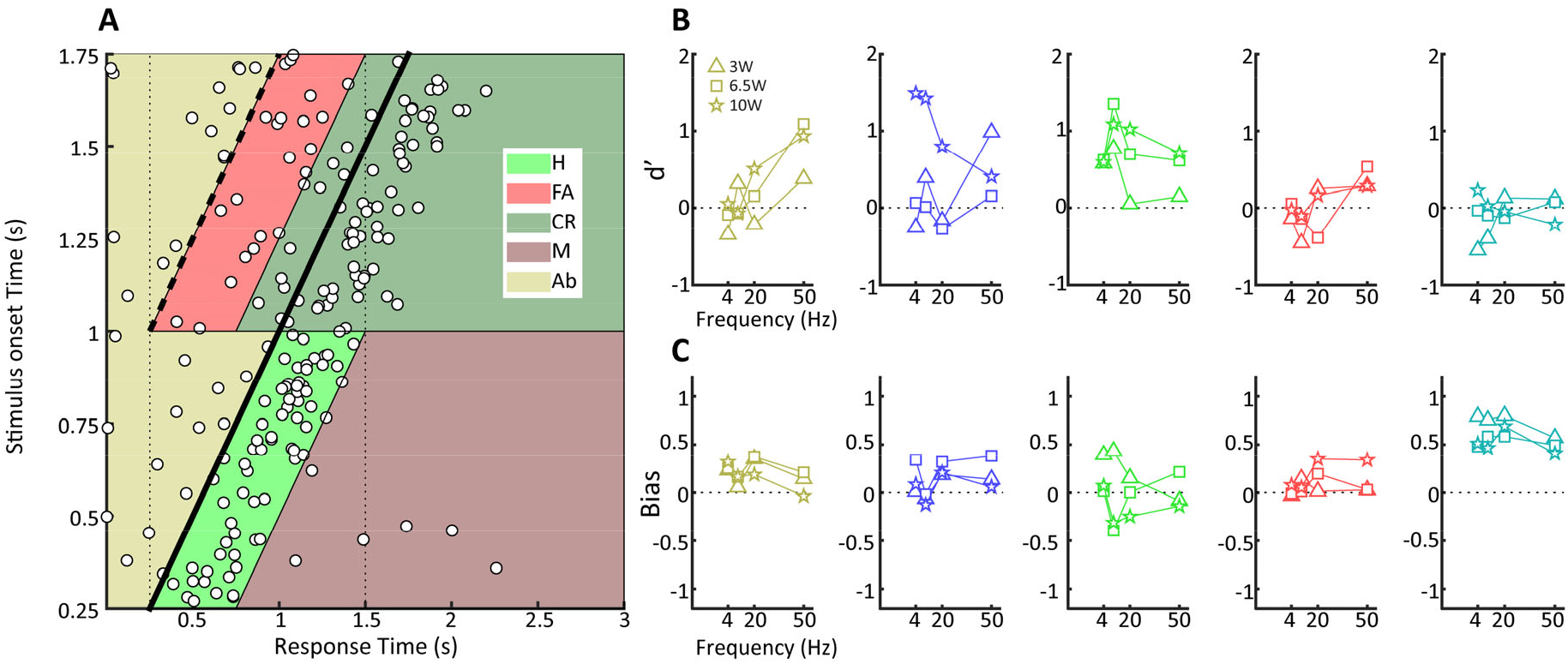
Behavioral detection of optogenetic activation of LGN CaMKII neurons. (A) As Figure 2B, but for an animal detecting blue LED activation of CaMKIla neurons in the LGN. Note that while the animal can clearly perform the task, detection performance is more variable than for the visual stimulus. (B) d’ values at the three LED intensities used are plotted for each animal at four stimulation frequencies (4,10, 20 and 40 Hz). (C) Same as B but for Bias scores.

Successful utilization of a prosthesis requires not only detection of stimuli, but also necessitates discrimination between stimuli in terms of both spatial and temporal aspects. While spatial aspects related to coverage of the receptive field are much studied and discussed ^45^, temporal aspects are likely at least equally important, particularly considering the usefulness of dynamic stimulation paradigms ^13^. We therefore proceeded here to study limitations in discrimination of temporally modulated visual stimuli, i.e. flicker, to provide a basis for comparison to future optogenetic work. Here, we used operant schedules as these are particularly sensitive and resistant to behavioral extinction to assess the capacity of tree shrews to discriminate among various visual flicker frequencies. After preliminary behavioral shaping (see methods), tree shrews continuously nose poked for reward on a variable interval VI-30 schedule, with stable response rates of about 0.25Hz (Fig.5A). Animals were then transferred to the discrimination task, where responses in the presence of the 20 Hz target flicker were still rewarded on the same VI 30 schedule (S+), while nose pokes in the presence of other non-target flicker frequencies were not rewarded (S-). Animals received 8 days of training for each frequency discrimination. An example cumulative record of nose pokes during S- and S+ conditions for an animal that has acquired the task on the 8^th^ day of training for the initial 1 vs 20 Hz discrimination (Fig.5B). Note that nose pokes during the S-become rare, while the animal’s nose poking during the S+ continues unabated. Using signal detection theory (see methods), we then computed hit rate, false alarm rate and d’ sensitivity for the 8 consecutive days of discrimination training in the 1 vs 20 Hz condition (Fig.5C). With training, d’ sensitivity rises from 0.6 to 3.2 due to both reduced false alarm and enhanced hit rates. All tree shrews showed a general improvement in task performance over training days, as measured both by percent correct and d’ (Fig. 5D,E), reaching values between 60% and 90% correct corresponding to d’ sensitivities in the range of 1.0 to 3.5 (oneway ANOVA [F(6,42)=29.6], p<0.001 for % correct, [F(6,42)=9.3], p<0.001 for d’). Bias analyses reveals substantial variation among animals (Fig. 5F), with a general trend of reduced overall bias with training except for one tree shrew (cyan trace) that achieved modest d’ sensitivity due to elevated false alarm rates, i.e. high positive bias values. Overall response rates in the S+ and S-conditions (Fig.5G) decreased and increased over training days in the S- and S+ respectively (oneway ANOVA: S+ [F(6,42)=3.1], p<0.05, S- [F(6,42)=8.0], p<0.001). Interestingly, rats performing a discrimination on a VI-30 reward schedule, only show a significant decrease in the S-response rate, with no concomitant increase in response rates during the S+ ^46^, revealing a potentially interesting inter-species difference. Following acquisition of the 1 vs 20Hz task, the animals spent 8 days of training on each of the remaining discrimination tasks to test transfer of frequency discrimination capacity and determine associated thresholds (see methods). All tree shrews readily transferred between the 1Hz vs. 20Hz and 5Hz vs. 20Hz conditions (Fig. 6A). As the non-target frequency approached the target frequency, d’ sensitivity declined. All animals had trouble transferring to the 10Hz vs. 20Hz condition, although one animal (black trace) was able to relearn this problem. All tree shrews had trouble acquiring the 30Hz vs. 20Hz condition, but 40Hz vs. 20Hz was easier to acquire with one animal (green trace) even rebounding to about 1.5 d’ values suggesting transfer from previously learned conditions. The average d’ sensitivity for n=6 tree shrews that completed this study (Fig.6B) suggests that for a standard frequency of 20Hz, comparison frequencies must be about 15Hz away to be easily discriminated by tree shrews. For frequency pairs spaced closer together, response latency was similar for S+ and S-trials (Fig.6C), and interestingly response rate for the 20Hz stimulation did not remain invariant but increased with separation of frequency difference between the two frequencies tested (Fig.6D). Having examined the limits of tree shrew discrimination performance for visual flicker as described above, we wanted to explore LGN and V1 neural activations to this type of stimulus and estimate decoding performance of single neurons in terms of flicker discrimination.

**Figure 5.**
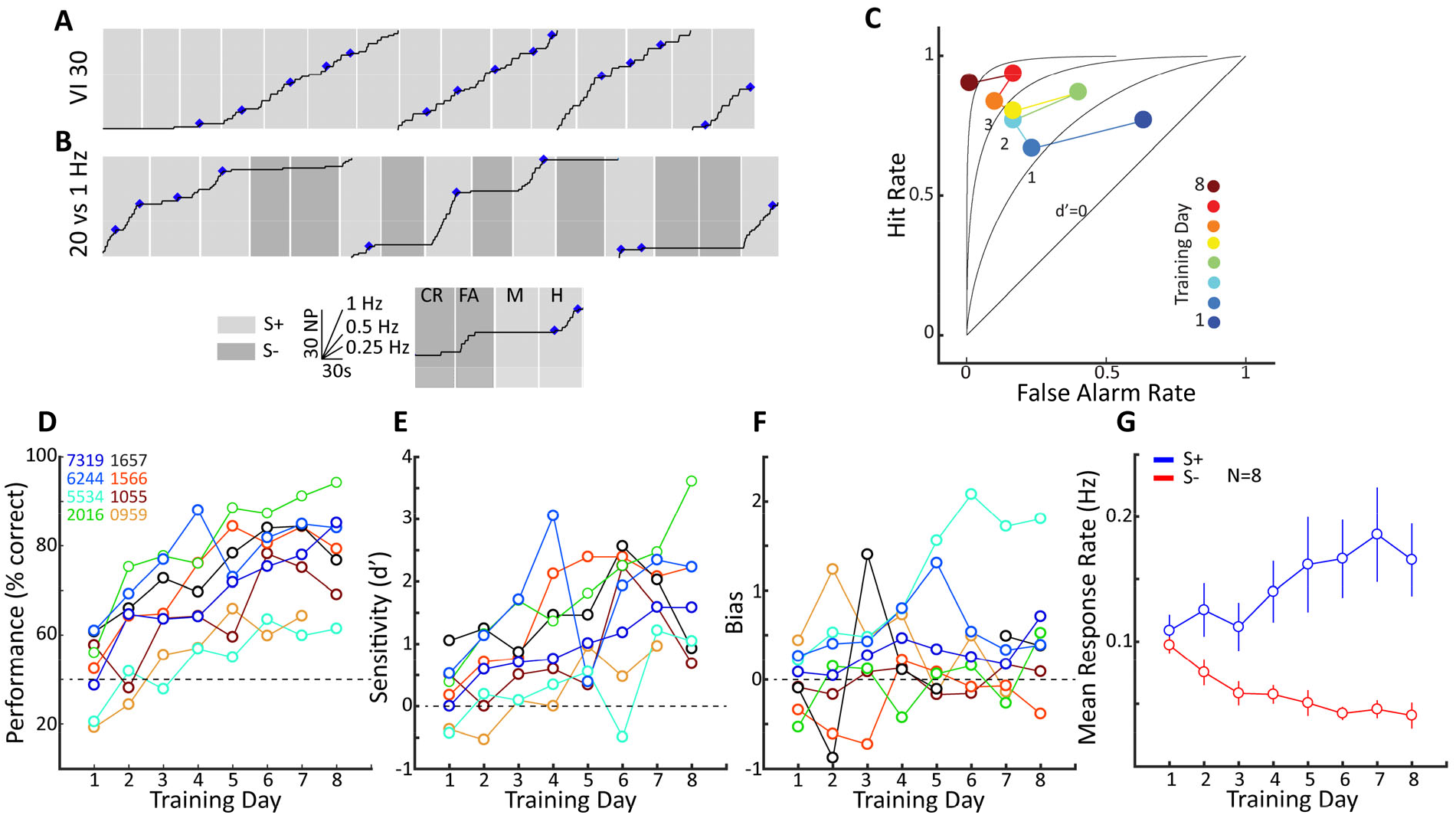
Flicker frequency discrimination in tree shrews. (A) An example cumulative record of tree shrew nose poking under a VI3O schedule of reinforcement, showing the typical steady response rates under this schedule. Each rectangle corresponds to a trial, and each upward tick corresponds to a single nose poke with the and the abscissa being time. Thus steeper slopes reflect higher nose poke rates, see inset. The record resets to zero after 60 nose pokes. (B) A cumulative record for the same animal following acquisition of a 1 vs 20 Hz flicker discrimination task. Light and dark shading of the rectangles represent S+ and S-trials respectively. Note that following learning, nose pokes in the S-become rare, while nose pokes in the S+ show higher rates than during the VI 30 schedule in A. In order to calculate d’ we defined correct rejections (CR), false alarms (FA), misses (M), and hits (H) as exemplified in the inset. (C) False alarm vs. hit rate for the same animal as in A and B, across learning days illustrating a steady increase in sensitivity over training sessions. (D) Performance over training days on the same 1 vs 20 Hz flicker discrimination task for all animals as measured by percent correct. Colored numbers in the inset reflect individual tree shrews. (E) Same as D but with performance measured using d’. (F) Bias scores over training days for the same animals. (G) Mean response rate is plotted over training days for nose pokes in the S+ and S-. Note the steady increase in response rates during the S+ trials with a concomitant decrease in response rates during the S-trials.

**Figure 6.**
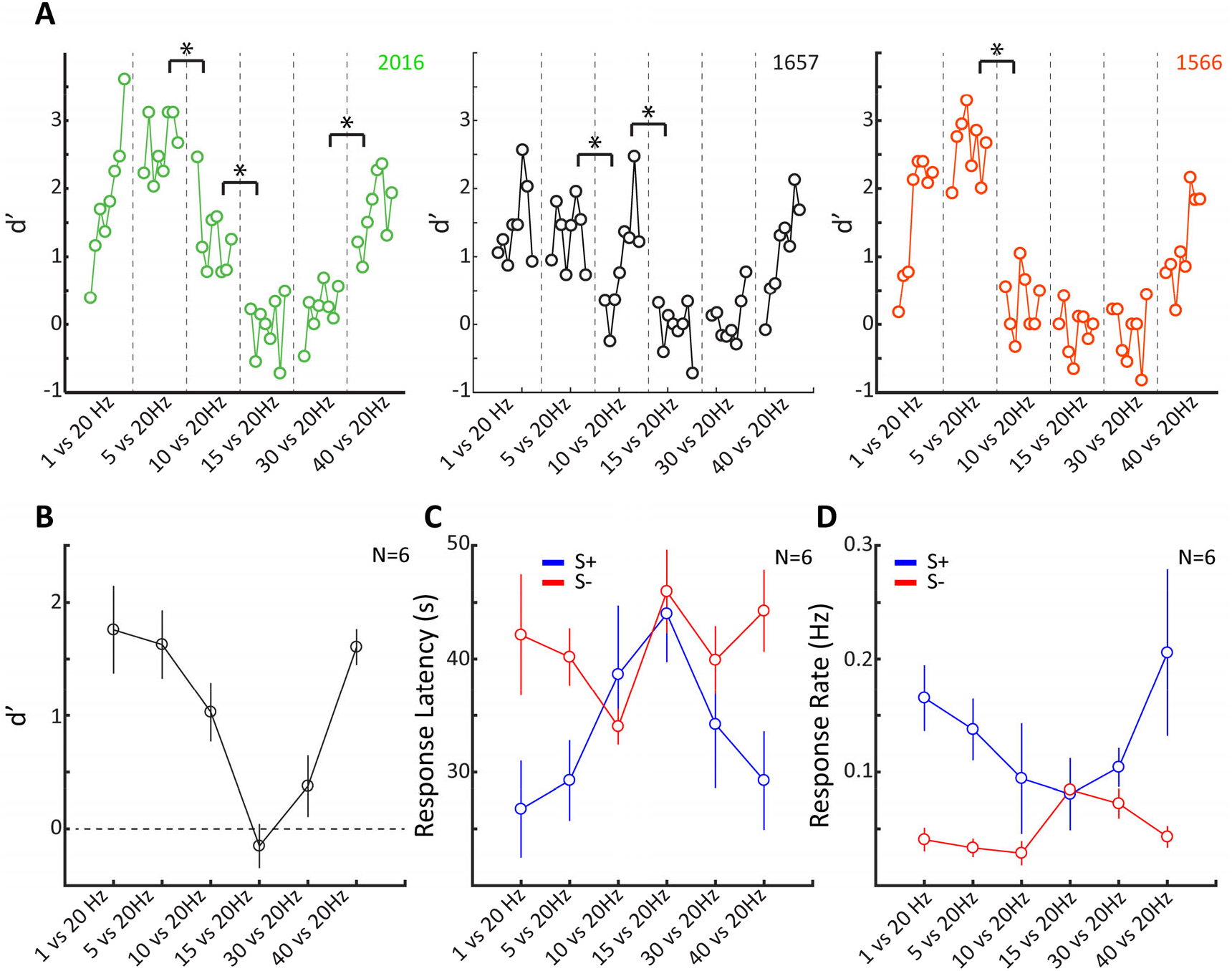
Perceptual limits on visual frequency discrimination in tree shrews. (A) Tree shrews were trained for 8 days on 6 different discrimination conditions in succession.The S+ was always 20 Hz, while the S-first increased in frequency from 1 to 40 Hz. d’values are plotted for 3 individual animals as training progressed under each successive condition. Significant changes in performance between the last 3 sessions of the previous discrimination condition, and the first 3 sessions of the subsequent condition are indicated. (B) Same as A, but mean values for all 6 animals, and only for the final sessions of training under each condition. In all figures error bars reflect +/- SEM. (C) Response latencies from the onset of the flickering stimulus until the first nose poke during S+ and S-trials on the final day of training for the different discrimination conditions for all animals. (D) Same as C but for response rates during the S+ and S-conditions.

Understanding the relation between perception and neural activations is of central importance for improving future generations of visual prosthetics. We focus here on activity from isolated LGN and V1 neurons, using full field flickering visual stimulation that covered the receptive field of each neuron under study delivered at 10, 20 and 40Hz (see methods). Fig.7A shows spiking responses of a single LGN neuron and an example single trial Fourier transform for each flicker frequency. Note that peaks are generally related to the flicker frequency as well as harmonics at integer multiples. On the right panels, single trial Fourier peak amplitudes are plotted, with each panel corresponding to a different flicker frequency. We used signal detection theory to estimate d’ sensitivity and bias for each neuron, in terms of its sensitivity for discrimination between the target visual stimulation frequency 20Hz and the two non-target frequencies 10 and 40Hz (see methods). For 10Hz visual flicker, the peak at 10Hz always exceeds the peak at 20Hz, such that all trials are classified as “correct rejections”. For 20Hz flicker, the 20Hz peak is larger than 10Hz peak for all trials, corresponding to “hit” responses. Accordingly, this LGN neuron is highly sensitive (d’=3.3) to both the 10Hz vs 20Hz and 40Hz vs 20Hz discriminations. A scatterplot reveals that indeed a substantial fraction of LGN neurons were sensitive to both discriminations (Fig.7B), corresponding to a cluster of neurons in the upper right part of the diagram. The LGN d’ values were accordingly more skewed towards high sensitivity (Fig.7C), while bias was significantly different between the two discriminations (Fig.7D), with preponderance of “miss” errors on the 10Hz vs 20Hz discrimination and “false alarm” errors on the 40Hz vs 20Hz discrimination; an effect we attribute to the harmonics of the Fourier transform. Repeating this analysis for a population of V1 neurons, we found a strikingly distinct activity pattern. Fig.8A shows spiking responses of a single V1 neuron and an example single trial Fourier transform for each flicker frequency. For 10Hz flicker, the 1^st^ harmonic peak at 20Hz actually exceeds the 10Hz peak on many trials, so that these trials are designated as “false alarms”. For 20Hz flicker, the 20Hz peak is larger than 10Hz peak for all but one trials, corresponding to largely “hit” responses. Accordingly, this V1 neuron is insensitive (d’=0) to the 10Hz vs 20Hz discrimination. However, it is highly sensitive to the 40Hz vs. 20Hz discrimination, with a d’=2.9. Other V1 neurons (Fig. 7B) exhibited the opposite pattern, with high d’ sensitivity observed for the 10H vs. 20Hz discrimination. A scatterplot of d’ values (Fig.7C) highlights that distinct populations of V1 neurons carried information about each of the two flicker discriminations, with few neurons carrying information for both. The distributions of d’ values for both discriminations were relatively broad (Fig.7D), suggesting that flicker-related information is dispersed among the V1 neural population, with similar mean values around d’=1.8. Bias distributions were centered around zero (Fig.7E) and also quite broad, as many neurons were only weakly sensitive to the flicker discrimination. Comparing LGN and V1 d’ sensitivity, we found that in V1 there is a substantially smaller group of neurons with high d’ values (d’>2.0) for both discriminations (n=13) compared to high d’ for only one or the other individual discriminations (n=29). The difference to corresponding numbers in the LGN (d’_both_: n=25, d’_single_: n=18) is significant (χ^2^ test, p<0.05). This certainly suggest that both LGN and V1 contain substantial populations of neurons whose activity provides sensitive information about visual flicker. While LGN neurons appear to multiplex information, i.e. high d’ for both discriminations, the information in V1 is distributed to more specialized neurons that can solve one or the other problems but not both. The spreading out of information across the V1 neural population may pose a challenge for downstream brain areas, which need to utilized inputs from distinct populations to solve the different discriminations. Our neural recordings thus provide novel insights into how the neural representation of temporally structured visual information is transformed from LGN to V1.

**Figure 7.**
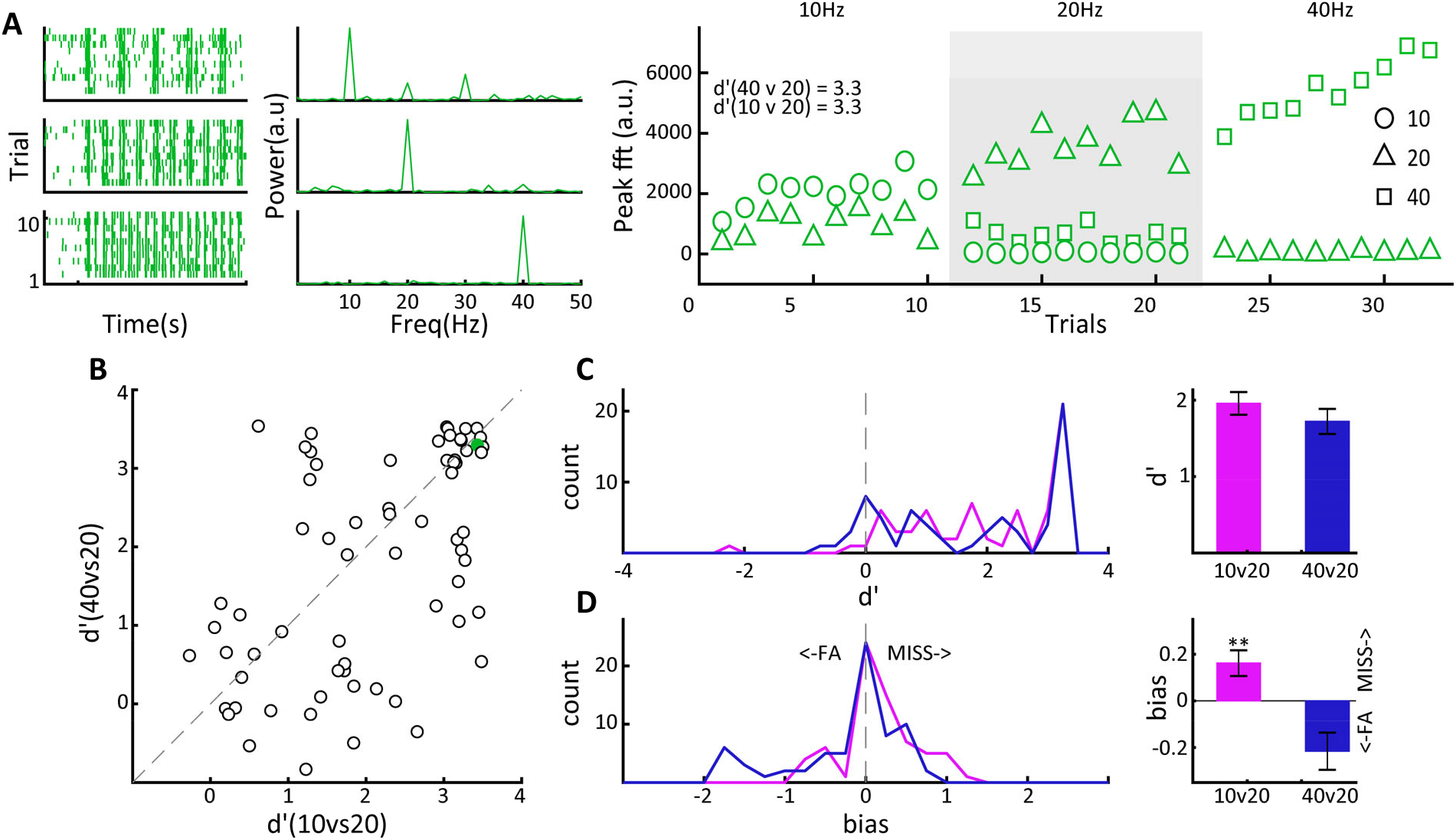
Flicker frequency discrimination is multiplexed in LGN neurons. (A) Responses of an example LGN neuron that discriminates both 40 from 20 Hz as well as 10 from 20 Hz. Responses to 3 different frequencies of full field flicker are displayed in raster plots (left) and their associated FFTs (center). The right panel shows the relative peaks in FFT power over 10 trials for the three frequencies of flicker stimulation. (B) Scatter plot of d’values calculated for the 10 vs 20 Hz vs the 40 vs 20 Hz discrimination conditions. The green dot represent the neuron plotted in A. Note that a small jitter has been added in order to make overlapping data points visible. (C) Histograms of d’values for all LGN neurons recorded for the 2 discrimination conditions are shown at left, with mean d’values for the 2 conditions shown at right. (D) as D but for the bias values calculated from the 2 discrimination conditions.

**Figure 8.**
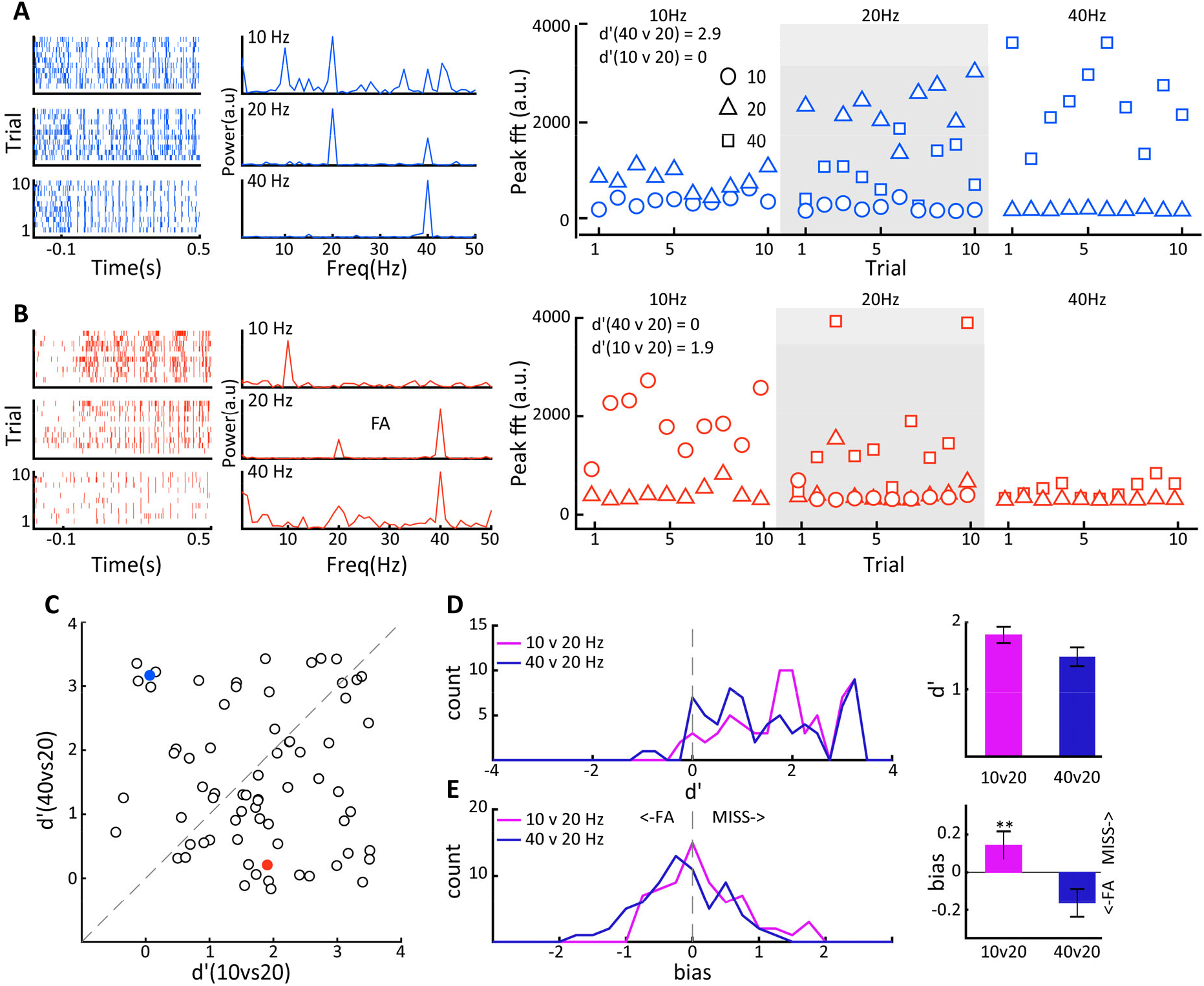
Flicker frequency discrimination is distributed in VI neurons. (A) Responses of an example V1 neuron that discriminates 40 from 20 Hz but not 10 from 20 Hz. Responses to 3 different frequencies of full field flicker are displayed in raster plots (left) and their associated FFTs (center). Note that for the 10Hz flicker, the harmonic at 20 Hz is actually larger than the fundamental at 10 Hz. The right panel shows the relative peaks in FFT power over 10 trials for the three frequencies of flicker stimulation. (B) is as A, but for a neuron that discriminates 10 from 20 Hz but not 40 from 20 Hz. (C) Scatter plot of d’values calculated for the 10 vs 20 Hz vs the 40 vs 20 Hz discrimination conditions. Blue and red dots represent the neurons plotted in A and B respectively. Note that a small jitter has been added in order to make overlapping data points visible. (D) Histograms of d’values for all VI neurons recorded for the 2 discrimination conditions are shown at left, with mean d’values for the 2 conditions shown at right. (E) as D but for the bias values calculated from the 2 discrimination conditions.

## Discussion

We demonstrate here that optogenetic activation of the visual thalamic LGN triggers local neuronal circuit activity as well as an associated artificial visual percept that can be readily detected by the animals. A previous report has already provided proof of principle that electrical LGN stimulation can elicit artificial visual percepts in macaque monkey ^47^. This previous study used a visually guided saccade task instead of the nose-poke detection paradigm in our study, but essentially both tasks permit the characterization of behavioral responses to natural and artificial visual percepts across the visual field. As macaques in the previous study, tree shrews immediately generalized from the visual to the optogenetic task, suggestive of a visual nature of the evoked artificial percept. The overall task performance we observed for optogenetic stimulation reached maximum d’ values of around 1.5 and thus remained somewhat lower than the 2.5 ceiling values observed during visual stimulation. However, the animal that achieved best d’ during optogenetics actually performed comparatively poorly on the visual detection task (dark blue traces Fig.3, 4), suggesting that detection of natural and artificial visual stimulation does necessitate somewhat divergent behavioral strategies and possibly attentional focus. Indeed, it has been shown that for macaques, extensive training on optogenetic stimulus detection in V1 is in fact detrimental to detection of visual stimuli ^48^. This finding emphasizes the potentially significant impact of learning, neural plasticity and behavioral strategy on optogenetic stimulus processing. Such aspects may also underlie the interindividual differences we observed to stimulation parameters, but variations in factors such as virus expression and LED probe placement might also contribute. Indeed, computational modeling work relating to V1 suggests that the spatiotemporal configuration of light delivery and virus transfected neural elements including dendrites plays a crucial role in how optogenetic activation is translated into cortical activity and ultimately perception ^49^. Further work extending computational modeling of electrical activations ^50^ to optogenetics in the LGN is necessary, ideally refining the models to integrate available experimental data. Our present results suggest that optogenetic LGN stimulation, up to the light intensities tested here, produces robust artificial percepts that tree shrews can readily detect in a suitable behavioral paradigm, although there are substantially more errors, mostly of the false alarm type, than in a comparable task involving natural visual stimulation. Increasing stimulation intensity further may be one option to enhance performance, as long as this does not cause tissue damage, nonspecific activation due to heating or undesirable loss of spatial selectivity ^51^. Note that overly strong LGN stimulation is in any case undesirable, as it has been shown to deactivate visual brain areas downstream of V1 ^52^, which would certainly compromise goal directed behaviors based on artificial visual percepts.

There are other more principled factors that might limit the detectability of optogenetic stimulation, notably the co-activation of ON and OFF pathways. These signals arrive in V1 layer 4, such that both pathways will be concurrently stimulated during electrical or optogenetic manipulations, unlike during natural vision where ON and OFF neurons are activated in alternation ^53,54^. Such coactivation will thus produce cortical activation patterns not encountered during natural vision, potentially explaining the perceptual performance decrement for artificial vision. In the tree shrew LGN, ON- and OFF-neurons tend to be segregated in separate layers, with layers 1,2 containing ON cells, layers 3-5 containing OFF cells and layer 6 containing a mixed population ^18^, note that we have adopted the conventional labelling the LGN layers as 1-6 from medial to lateral, whereas in Conway this ordering is reversed. Since we targeted medial LGN layers 1-3 in the present study (see Fig.1A), we likely also activated a mixed population here as our focus was obtaining proof of feasibility related to perception of LGN activation. Future experiments, targeting for example exclusively ON layers 3-5 of tree shrew LGN, can however address the issue of concurrent ON/OFF activation. We predict that such stimulation might trigger artificial percepts to which animals exhibit elevated d’ sensitivity values more similar to those for natural visual stimuli, and that detection performance under these conditions will be more sensitive to a visual masking stimuli containing dark structure rather than bright structure on a gray background. The functional anatomy of the tree shrew visual pathway permits a systematic investigation of this issue, providing an intermediate step before the issue can be addressed in macaques where LGN layers are also segregated for ON and OFF pathways ^55^, in a translational neuroscience context towards eventual human clinical application development.

As current findings in visual prosthetics point towards a crucial role of temporal patterning of artificial stimulation ^13^, we studied the perceptual capacity of tree shews in discrimination of temporally structured visual stimulation, i.e. flicker, as well as corresponding neural circuit activation in LGN and V1. Using variable interval operant schedule training, we showed that tree shrews can perform discriminations between distinct flicker frequencies with high sensitivity, achieving d’ values up to about 3.5. We suggest that operant schedule task performance may be more suitable to quantitatively examine perceptual abilities with high precision; note for example performance of the animal that achieved d’≈1.5 on the detection task but d’>3 on the discrimination task (Figs. 3D, 5E red traces). The learning curves (Fig.6A) document that tree shrews can learn to discriminate among sensory stimuli within a few days as well as generalize to novel stimuli. Further, in our experiments without food or water scheduling, animals remained consistently engaged throughout 200 trials of daily sessions in an unconstrained freely moving behavioral context. This underscores the visual perceptual capacities of the tree shrew, also documented by other studies ^28,31,32,56^ in terms of translational vision neuroscience.

Making progress in visual prosthetics necessitates a better understanding of how task-relevant visual signals are transformed as they ascend the visual pathways and how they relate to behavioral performance. In our LGN and V1 recordings to visual flicker at the different flicker frequencies (10, 20, 40Hz), we therefore aimed to quantify how much information about two discriminations between pairs of frequencies (10 vs. 20, 40 vs. 20Hz) were contained spike trains of LGN and V1 neurons. We used Fourier analysis in conjunction with signal detection theory, which allowed us to compute a d’ sensitivity that could be readily compared to the d’ values obtained in the behavioral studies. Similar to previous studies in several mammalian species ^57–59^ including the tree shrew ^28^, we found robust entrainment of spiking activity in the employed 10 - 40Hz frequency range. Examining d’ sensitivity for the two discriminations, we found that many LGN neurons could solve both discriminations (Fig.7B), while V1 neurons tended to possess information about one or the other but not both discriminations (Fig.8C). This result is compatible with previous findings showing that LGN neurons are much more stereotyped than V1 neurons in terms of their frequency response, exhibiting cutoff values above 30Hz ^57^. By contrast, V1 neuron frequency preference was much more distributed along the 1-40Hz continuum. This implies that LGN neurons can follow visual flicker at 40Hz and lower frequencies, whereas V1 neurons have bandpass characteristics and only respond to particular frequencies. Crucially, the information contained in the LGN spikes is not lost, but transformed from a multiplexed system where single LGN neurons carry aggregate information, to an explicit representation in V1 where single neurons convey information allowing discrimination between particular temporal frequencies. The apparent lack of loss between LGN and V1 is consistent with previous findings that examined spiking variability across these brain regions ^60^ and d’ sensitivity estimates for LGN and V1 obtained in macaques to drifting sinusoidal gratings ^61^. It is established that visual signals are transformed in various ways as they pass from the LGN to V1, leading to emergent properties such as orientation tuning or integration of ON and OFF pathways into a complex spatiotemporal sensitivity profile ^62,63^. These V1 activity properties make information implicit in the LGN relay cells explicit across the cortical neuronal population. We suggest that the same is occurring for temporal frequency. The transformation of information representation from LGN to V1 has important implications for potential visual prostheses. In particular, temporal aspects of LGN neural activation will be crucial in determining which population of downstream V1 neurons are activated by the same set of relay cells. We suggest that these effects of temporal patterning may be considerably larger when LGN is activated compared to V1, which may be advantageous for artificial vision as more information can be transmitted by a given number of light sources. The characteristics of the V1 neural population response elicited in artificial vision will clearly have a profound impact on downstream cortical visual area responses in extrastriate and inferior temporal visual areas, which underlie perception and memory for objects and other behaviorally relevant stimuli. At present, it seems likely that artificial stimulation strategies that most closely emulate neural activations occurring in natural vision are most suitable to be readily interpreted by higher visual areas. This would permit an intuitive form of artificial vision that does not require extensive learning-dependent restructuring of inter-area connectivity inside the visual cortex. Nevertheless, the nature of neural circuit activations in higher visual areas triggered by artificial vision will need to be explored further to optimize artificial stimulation strategiesm and ultimately also acceptance of such prosthetic devices by individuals with impaired vision.

In summary, our findings reinforce the idea that the LGN might be a particularly promising target for artificial vision ^54^. Our findings suggest that the tree shrew, as a small diurnal highly visual mammalian species, appears well suited for translational animal work in the area of visual prosthetics before the validated approaches are tested in macaques and eventually humans. We have started here with optogenetics using the CamKIIα promotor and shown that tree shrews can readily detect single channel light stimulation delivered within the LGN. Expanding the scope to additional promoters, for example targeting other classes of thalamocortical relay cells, as well as multiple light sources that allow dynamic, sequential activation of neural circuits ^64^ are highly promising avenues towards a future generation of prosthetic devices. For example, intermittent optogenetic inhibition of LGN inhibitory cells could provide an efficient way to trigger coordinated activation in nearby relay neurons. Our findings show robust expression of light-sensitive opsins in layer 3 as well as layer 4 of cortex, opening the possibility of illuminating the thalamocortical axon terminals by light arrays overlying the cortical surface. Such devices can be manufactured as flexible implants, which could eventually deliver stimulation in a significantly less invasive manner compared to implanted electrodes, highlighting a potential advantage of the LGN as a target for artificial vision.

## Materials and methods

### Subjects

Male and female tree shrews (Tupaia Belangeri) weighing 235±10 g (0.6-2 years old) participated in the experiments. They were given ad libitum access to food and water and housed in temperature-controlled rooms (ambient temperature: 26±1°C, air humidity: 60±5%) under a 13/11 light/dark cycle with gradual illumination transitions at the beginning and end of the light period. All experimental procedures were performed in accordance with applicable regulations and approved by the authorities.

### Behavioral training

Tree shrews were first shaped to enter a nose poke mounted on a transparent horizontal platform at the center of a 70cm diameter opaque sphere for mango juice reward using continuous reinforcement in daily sessions lasting 30 minutes in the absence of food or water restriction. The mango juice was delivered at a fluid port located on the edge of the platform behind the nose poke and adjacent to the sphere wall.

For the detection task, animals were trained to wait in the nose poke until a bright 10°x10° bright square was projected onto the globe directly ahead of the animal. The target onset times were gradually increased from fixed 250ms to fixed 250ms + randomized (0 to 1500 ms) duration, while concurrently the response window for obtaining a reward was decreased from 1200ms to 500ms. Once animals achieved good performance on this task, a cloud of bright moving dots 2° in diameter resembling a phosphene was used as a stimulus with unchanged timing parameters (fixed 250ms + randomized 0 to 1500 ms). Animals participated in seven training sessions with the visual stimulus in a fixed position (15° elevation, 30° right of center), see Fig.3A-D. For the visual field generalization task, the phosphene stimulus was horizontally displaced by 30° consecutively, up to a position of 150°. Behavioral sessions generally lasted 200 trials, taking approximately 30 to 60 minutes.

For the discrimination task, tree shrews progressed successive fixed ratio (FR) and variable interval (VI) schedules until animals exhibited stable responding on a VI-30 schedule, with all preliminary training taking place in the constant presence of a full field light flickering at 20 Hz. In FR-x training, animals received a reward after x nose pokes, and we proceed from FR-2 (30 min each session) to FR-4, FR-6, and FR-8 (60 min each session), which lasted for 4 days in total. We then switched to VI training, which included 3 phases (60 trials each session, 50-70 seconds per trial): VI-10, VI-20, and VI-30 lasting for 2 days per phase. On the final VI-30 schedule, a reward becomes available for a single lever press each time a random duration, 30±15s, has elapsed. Animals were then switched to the discrimination task, where trials of 20Hz flicker were maintained (S+) but trials with a comparison frequency flicker were randomly interspersed (S-), during which no reward was available. Animals were trained over 8 days on each of 6 successive discrimination tasks with different non-target S-frequencies (1,5,10,15,30, and 40 Hz), with S+ always remaining at 20 Hz.

### Behavioral data analysis

The use of signal detection theory ^65^ for our detection task data requires estimation of hits, false alarms, misses and correct rejections (see Fig.3A). We thus partitioned the trials into two halves depending on target onset time, each of 750ms duration. For onset times from 250 to 1000ms, responses that occurred after target onset and before the 500ms response window were assigned as “hits”, and responses occurring after the response window were assigned “misses”. We then constructed a time window identical to the “hit” window for the second half of the trials with onset times 1000 to 1750ms and used this to estimate “false alarms”. In terms of timing, “hit” and “false alarm” windows are indistinguishable to the animal, as these windows cover identical time periods, the only difference being that no target occurs in the “false alarm” window. If animals do not respond during the “false alarm” window, the trial is assigned as “correct rejection”. This procedure permits the computation of sensitivity (d’) and bias (b) parameters according to signal detection theory for response times generated in a simple detection task. Responses occurring prior to the “hit” and by extension “false alarm” windows were considered as aborted trials. Rates were computed as follows: 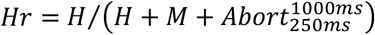, 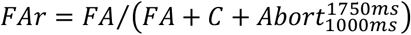. Sensitivity: *d′* = *z*(*Hr*) – *z*(*FAr*), Bias: 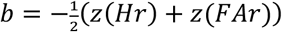. Excellent behavioral performance on decision tasks is associated with high sensitivity and low bias.

For the discrimination task, to determine the discriminability of the different stimuli according to signal detection theory, we calculated d’ sensitivity as follows: β or more responses represented a hit (H) or false alarm (FA) on S+ and S-trials respectively, while less than β responses assigned as miss (M) or correct rejection (CR) on S+ and S- trials respectively. The value of β as an arbitrary threshold was chosen as 6 nose pokes in the present analysis. To assess transfer of discrimination capacity to the subsequent non-target frequency, we compared d’ values on the last 3 days training in of the previous condition to d’ values on the first 3 days of training on the subsequent condition. A significant difference indicates lack of transfer between the two task conditions.

### Surgical procedures

For both viral injections/LED stimulator implantation and neural recordings, general anesthesia was induced by Alfaxalone (40 mg/kg, half dose in each leg, i.m.)^66^ and we administered Atropine (0.08 mg/kg, i.m.) to reduce secretions and Baytril (2.5%, 0.3 ml/kg, s.c.) to prevent infection. Anesthesia was maintained by isoflurane (0.5-3.5%) in 100% O_2_ administered using a nose cone for viral injections or endotracheal intubation for neural recordings. Here, after identifying the vocal cords, a PE50 polyethylene tubing extending 1cm from an endotracheal tube (Original Perfusor type: IV-standard-PVC, 6 cm, 1.5 x 2.7 mm) was inserted into the trachea under visual inspection using video laryngoscopy ^67^. We monitored exhaled CO_2_ (Physiosuite, Kent Scientific) and ventilated animals at 100 bpm (Small animal ventilator 683, Harvard Apparatus). Animals were placed in a stereotactic apparatus, skin was shaved, periosteum retracted, cranial bone cleaned (3% H_2_O_2_) and craniotomies were drilled to for access to injection or neural recording sites under appropriate local analgesia (Lidocaine, 1%). Post-operative analgesia was administered before the end of surgical intervention (Buprenorphine, 0.05mg/kg, s.c.). We injected the construct AAV2-CamKIIa-hChR2(H134R)-mCherry (1 μl, UNC Vector Core) unilaterally using a microsyringe (34 GA. beveled Needle, 10 μ l NANOFIL syringe, WPI) into the left LGN (AP 3.1, ML 4.8, DV from 5-7 dependent on the neural response for 5 Hz full-field flickering stimulus to ensure accurate placement). Following viral injection, a wirelessly powered device containing the μ-LED at the tip end of a freely adjustable needle (NeuroLux, Chicago IL) was lowered into the LGN, again under stereotactic guidance, and cemented in place with dental acrylic (Paladur). After viral injections/LED stimulator implantation, the skin was sutured and animals were allowed to recover for 14 days before participating in further behavioral training. We allowed 4-6 weeks for viral expression before commencing optogenetic experiments. Neural recordings were performed using resin coated tungsten electrodes (Impedance ~300kΩ, FHC, Bowdoin ME) in LGN or V1, using visual stimulation delivered on a video monitor (VPixx, Canada), placed 28.5 cm from the animal and covering the visual field location corresponding to the receptive field of the neurons being recorded, as described in more detail previously ^68^. For optogenetic activation of the LGN, we employed an optrode, i.e. a 100 μm diameter optic fiber coupled to an electrode for simultaneous monitoring of neural activity close to light stimulation site. The optic fiber was connected to a 473-nm blue laser (Changchun New Industries Optoelectronics, China). Laser intensity was set to 3.5mW using an optical power meter (PM 100D, Thorlabs Newton NJ). Stimulation was delivered using a duty cycle of 50% controlled by a pulse generator (Rigol, Beaverton OR).

### Neural recordings

We recorded from a total of 120 and 74 isolated single neurons in LGN and V1 respectively, presenting full field bright visual flicker on a dark background covering the neurons’ receptive field and surrounding parts of the visual field. We used flicker frequencies of 10, 20 and 40Hz, presented for 1s duration with 10 repetitions per condition. We selected populations of responsive neurons (LGN: 68, V1:69), which showed significant enhancement of activity for any flicker frequency compared to baseline (paired t-test pre-stimulus baseline vs. flicker evoked activity, p<0.05) as well as mean spike rate exceeding 10 spikes/s. For single trial estimates of neural frequency response, we computed the Fourier transform of the spike train (FFT) binned at 1ms, and estimated the amplitude at frequencies 10, 20 and 40Hz. We then used signal detection theory to estimate d’ sensitivity, with 20Hz always serving as reference or target frequency in similarity to the behavioral experiments, and either 10Hz or 40Hz as non-target comparison frequency. Taking the example of 20Hz vs. 10Hz discrimination, a “hit” was assigned when 20Hz visual flicker was presented and the FFT peak at 20Hz was larger than the peak at 10Hz. If the 10Hz FFT peak larger, the trial was counted as “miss”. The “correct rejections” and “false alarms” were assigned depending on whether the 10Hz FFT peak was smaller or larger than the 20Hz FFT peak during 10Hz visual flicker presentation. This procedure allowed us to estimate d’ sensitivity and bias in an identical fashion as we did for the behavioral experiments using hit and false alarm rates computed as follows: *Hr* = *H*/(*H* + *M*), *FAr* = *FA*/(*FA* + *CR*). Sensitivity: *d′* = *z*(*Hr*) – *z*(*FAr*), Bias: 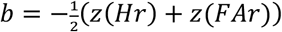. We do not wish to imply that the downstream brain areas are computing Fourier transforms, but use this procedure to assess the information present in single trial spike trains. To estimate how many bursts were elicited by LGN optogenetic activation, we used standard criterion of 50ms absence of activity followed by 4ms or less interspike interval between two subsequent action potentials.

